# Defining the minimal structural requirements of DivIVA in filamentous Actinomycetota

**DOI:** 10.1101/2025.06.24.661030

**Authors:** Maarten Lubbers, Belmin Bajramović, Véronique Ongenae, Joost Willemse, Dieuwertje de Bruin, Niels Mulder, Bastienne Vriesendorp, Francisco Barona-Gómez, Ariane Briegel, Gilles P. van Wezel, Klas Flärdh, Dennis Claessen

## Abstract

The morphogenetic protein DivIVA exhibits diverse functions across bacterial phyla. In Bacillota, DivIVA is primarily involved in cell division, whereas in Actinomycetota, it plays a central role in coordinating polar growth. Due to its essential nature, gaining insight into DivIVA function is challenging. Here we report on the functionality of truncated DivIVA proteins, using a unique *divIVA* deletion mutant created in cell wall-deficient *Kitasatospora viridifaciens* L-forms. DivIVA comprises an N-terminal domain, two coiled-coil regions separated by an intercoil linker, and a C-terminal domain. Deleting either the intercoil or the C-terminal region impacted branching dynamics. We also created a minimized variant wherein both were deleted simultaneously, containing the N-terminus and fused coiled-coils, resembling DivIVA from unicellular bacteria. Expression of this minimized variant resulted in severe growth defects. Cells exhibited a strong increase in hyphal width and cell wall thickness, accompanied by frequent tip bursting. Finally, we successfully introduced chimeric DivIVA from the unicellular actinobacterium *Mycolicibacterium smegmatis* with an N-terminal domain of *Kitasatospora viridifaciens*, demonstrating functional conservation within the phylum. In contrast, a chimeric DivIVA from *Bacillus subtilis* could not support growth, underscoring that polar growth is encoded within Actinomycetota-specific amino acid motifs encoded in the first and second coiled-coil. These findings enhance our understanding of the structure-function relationship for DivIVA and present new opportunities to study polar growth.

**Impact:** DivIVA is essential for polar growth in Actinomycetota. In Streptomycetaceae, this membrane-binding protein localizes at growing hyphal tips and along lateral hyphal walls where new branches emerge. Due to its essentiality, the structural relationship of DivIVA between unicellular and multicellular species remains elusive. Using a *Kitasatospora viridifaciens* L-form *divIVA* deletion mutant, we expressed truncated DivIVA variants to identify essential regions. Deleting two large unstructured domains produced a minimized variant containing the N-terminus and fused coiled-coils, resembling DivIVA from unicellular bacteria. This strongly impacted morphogenesis, increasing hyphal width and cell wall thickness, and leading to hyphal tip bursting. Finally, we successfully substituted DivIVA of *K. viridifaciens* with that from *Mycolicibacterium smegmatis*. Furthermore, *Bacillus subtilis* DivIVA could not facilitate reversion, showing that polar growth depends on amino acid motifs unique to Actinomycetota. These findings enhance our understanding of the structure-function relationship of DivIVA and offer new opportunities to study polar growth.

## Introduction

Streptomycetaceae are filamentous bacteria that form multicellular networks. Unlike well-studied bacteria such as *Escherichia coli*, which grow by adding new cell material along their lateral walls (1), *Kitasatospora* forms so-called hyphae that grow by extension at their tips. During this process, known as polar growth, new material is added only at the ends of the filaments (2). These filaments can also branch, leading to the establishment of a network of interconnected hyphae, called a mycelium. Polar growth and the formation of branches are orchestrated by DivIVA (2), a membrane-binding protein found at growing tips (3). DivIVA is an essential protein, and partially depleting DivIVA results in the formation of abnormally shaped hyphae and irregular branching (4, 5). In contrast, overexpressing DivIVA causes peptidoglycan to be added at multiple new sites, triggering excessive lateral branching (3, 5). In other unicellular Actinomycetota (or Actinobacteria), such as *Mycolicibacterium smegmatis* and *Corynebacterium glutamicum*, DivIVA is also essential, where it plays a central role in coordinating polar growth (6-8).

In Streptomycetaceae, DivIVA consists of five domains: an N-terminal domain, followed by a smaller coiled-coil, an intercoil region, a larger coiled-coil, and a C-terminal domain (5). The N-terminal domain and the two coiled-coils are highly conserved among Streptomycetaceae, in contrast to the variable intercoil region and C-terminal domain (5). DivIVA associates with negatively curved membranes, possibly through molecular bridging of the curvature by DivIVA multimers (9). Computer simulations predicted that the N-terminal coiled-coil domain binds to the membrane, with its affinity dependent on membrane lipid composition (10). Furthermore, DivIVA forms a multimeric complex in streptomycetes and long filaments *in vitro* (4). In non-polar growing bacteria, such as Bacillota (or Firmicutes), DivIVA is non-essential, while still playing a role in cell division. In *Bacillus subtilis*, the DivIVA protein targets the division septum and cell poles and contributes to the control of the topological specificity of cell division (11). DivIVA also plays a role in sporulating cells, interacting with the chromosome segregation machinery to aid in positioning the *oriC* region at the cell pole (12). Structural models based on crystallography indicate that the N-terminal membrane-binding domain of *B. subtilis* DivIVA forms a dimer with crossed-over loops, likely interacting with the membrane surface (13). Further, models based on the crystal structures suggest that the C-terminal coiled-coil region is involved in oligomerization and forms a tetramer (13).

A full understanding of the roles and mechanisms of DivIVA in Actinomycetota requires the ability to delete and functionally analyze this gene. This is made possible by using L-forms, bacterial cells that can proliferate without a cell wall and cell division machinery (14). L-form proliferation is governed by biophysical principles, driven by increased membrane synthesis, leading to an imbalance in the cell’s surface-to-volume ratio (15, 16). Previously, an L-form strain of *Kitasatospora viridifaciens* was generated under laboratory conditions and named *alpha* (17). *Alpha* only proliferates in its cell wall-deficient state when grown in osmoprotective media. When grown on medium with lower levels of osmolytes, *alpha* reverts to walled growth and forms mycelial networks comparable to the parental strain (17). The Δ*divIVA* was generated by deleting the *divIVA* gene, disabling the strain from reverting to walled, filamentous growth. Reintroduction of the native *divIVA* gene fully restored walled growth, showing efficient complementation *in trans* (18). Using this system, we investigated which domains of DivIVA are critical for its function in steering polar growth in Streptomycetaceae. For this, we used Δ*divIVA* to test which domains of DivIVA are required for walled growth. Therefore, we created a functional minimized DivIVA variant comprising the N-terminus and two fused coiled-coil domains. Amongst others, removing both the intercoil region and the C-terminal domain led to a very strong phenotype, with increasing cross-wall formation, highly irregular hyphal diameter, increased cell-wall thickness, and frequent lytic events. Furthermore, this minimized DivIVA protein structurally resembles DivIVA from unicellular bacteria, giving us a better understanding of structure-function relationships. This approach establishes a powerful platform for dissecting the functional architecture of essential morphogenetic proteins and enables novel strategies for engineering cell shape and growth in filamentous Actinomycetota.

## Materials and Methods

### Strains and media

Bacterial strains used in this study are listed in Table S1. *Escherichia coli* was grown at 30°C or 37°C at 200 RPM in LB. *Kitasatospora viridifaciens* DSM40239 was grown on solid MYM medium for 3 days (19) or as a liquid-shaken culture in TSBS (tryptone soy broth with 10% sucrose) at 200 RPM (20). L-form strains were grown on LPMA plates supplemented with 5% (v/v) horse serum and 25 mM MgCl_2_ (17), whereas L-phase broth (LPB) (17) was used for growth in liquid (at 100 RPM). To facilitate the reversion of the Δ*divIVA* mutant carrying different *divIVA* plasmids, strains were first grown in LPB medium for two days at 30°C at 100 RPM until reaching an OD of 0.5. 200 µL of these cultures were inoculated on reversion plates (MYM:LPMA (4:1)) plates and grown at 30°C for three days (21).

### Plasmid construction and transformation

Plasmids used in this study are listed in Table S2. All primers are listed in Table S3. To isolate genomic DNA of *K. viridifaciens*, the protocol was performed as described by Kieser et al. (20). Fragments were amplified using Q5 DNA polymerase (New English Biolabs). For Gibson assembly, pIJ82 containing the *gapA* promoter (22) was used as the backbone vector following digestion with *Xba*I and *Nde*I. Gibson-assembled products were transformed into competent *E. coli* TOP10 cells. Plasmids were isolated using the Nucleospin® plasmid Easypure kit (BIOKÉ) and confirmed by Sanger sequencing (Macrogen). Codon-optimized *divIVA* genes from *Bacillus subtilis* and *Mycolicibacterium smegmatis* were synthesized by Integrated DNA Technologies and Twist Bioscience, respectively. Plasmid transformation into the *K. viridifaciens* Δ*divIVA* mutant was carried out following the protocol described by Zhang et al. (18).

### Microscopy and imaging

#### Light microscopy

Microscopy analysis of mycelia grown in liquid was conducted using a Zeiss Axio Lab A1 upright Microscope, equipped with an Axiocam MRc (Zeiss) using a 40× magnification. Plate images were made using the Perfection V600 scanner (Epson). Colony photos were made using a Mikrocam SP 5.0 microscope camera (Bresser) connected to a SteREO Discovery V8 stereomicroscope (Zeiss).

To visualize growth over time, metal growth chambers shaped like a microscope slide with a 13 mm hole in the center were used. In these chambers, filtered MYM medium solidified with 1% agarose was enclosed between a Lumox Biofoil 25 membrane (Greiner Bio-One) and a cover slip (3). Mycelia were imaged at 30 °C under a Axio Observer.Z1 inverted light confocal microscope (Zeiss), using a Plan-Apochromat 40x/1.4 Ph2 objective and an ORCA Flash 4.0 LT camera (Hamamatsu).

To observe the bursting of hyphae in detail, liquid culture was spun down for 3 minutes at 0.4 rcf. To concentrate cells, the supernatant containing smaller mycelia was transferred to a new tube and spun down for 10 minutes at 13.000 rpm and resuspended 1:50 in filtered TSBS. A CellASIC B04A-03 microfluidic plate (Merck) was rinsed and primed with filtered TSBS. Further steps were followed as described by Passot et al. (23). An inverted Axio Observer.Z1 (Zeiss) was used with a 100X phase contrast objective.

#### Fluorescence microscopy

For DivIVA::eGFP localization studies, cells were imaged under an Axio Observer.Z1 inverted light microscope, using a Plan-Apochromat 100x/1.4 Oil Ph3 objective and an ORCA Flash 4.0 LT camera (Hamamatsu). L-form cells were placed on 1% agarose in P-buffer with 0.5 μg mL^-1^ FM4-64, while transformed *E. coli* cells were placed on 1% agarose in phosphate-buffered saline (PBS) in growth chambers. Metal growth chambers shaped as a microscope slide with a 13 mm hole in the center were used, where the medium was enclosed between a Lumox Biofoil 25 membrane (Greiner Bio-One) and a cover slip (3). For L-form strains, Z-stacks of images were acquired, with 25 planes with a 270 nm distance. To enable image deconvolution, point-spread functions (PSF) were determined using beads with excitation/emission wavelengths of 505/515 nm (green) and 633/660 nm (deep red) from the PS-Speck^TM^ Microscope Point Source Kit (Invitrogen). The beads were imaged and PSFs calculated using the Deconvolution module in Zen software (Zeiss, ver 3.9). Image stacks were captured with Z-spacing as above, and deconvolution was applied to the whole stack using the constrained iterative algorithm of Zen and the determined PSFs. Orthogonal projections were made from four focal planes through the center of the cells.

#### Scanning electron microscopy

To visualize colonies using scanning electron microscopy (SEM), agar disks (⍰ 8 mm) containing colonies grown on 80%MYM/20%LPMA plates were isolated using a hole puncher. The samples were prepared as described in the protocol of Yagüe et al. (24). The specimens were observed in a JSM-7600F field emission scanning electron microscope (JEOL) at 5.0 kV.

#### Transmission electron microscopy

For transmission electron microscopy (TEM), colonies grown on MYM/LPMA (80:20) plates were isolated. Samples were prepared as described by Yagüe et al. (24). The sliced samples were placed on copper TEM support grids and observed using a JEM-1400 Plus Electron Microscope (JEOL) at 80 kV.

#### Cryo-electron microscopy

For cell wall thickness measurements, sacculi were isolated and analyzed using cryo-electron tomography. 32 h old liquid cultures were resuspended in cold 0.1 M TrisHCl (pH 7.0), subsequently boiled in 4% sodium dodecyl sulfate (SDS) for 30⍰min, and washed with MilliQ. Samples were then enzymatically treated with DNase, RNase, and trypsin. After overnight incubation, the sample was again boiled for 30⍰min in 4% SDS and washed. Colloidal gold beads with a size of 10 nm were added to the samples in a 1:25 ratio, after which 3 µL was applied to a glow-discharged R2/2 200 mesh holey carbon EM grid (Quantifoil). The samples were vitrified in liquid ethane using an EM GP automated freeze-plunger (Leica) and observed using a single tilt specimen holder inside a 120 kV Talos L120C TEM (ThermoFisher) with a Lab6 electron source and Ceta detector at the Netherlands Center for Nanoscopy (NeCEN). Cell wall thickness was measured as described (25).

#### ImageJ analysis

To measure colony size, shape, and area, 30 colonies were picked per strain by randomly generating x- and y-coordinates and selecting the nearest free-lying colony to that point. Images were then measured with ImageJ version 1.53k by using a 5 mm reference image as a scale, after which the MeasureColony.js macro was run (26). Alternatively, the colony was selected with the magic wand tool or manually traced with the freehand selection tool. ‘Area’ and ‘Shape descriptor’ were automatically measured by ImageJ. Data was analyzed with R version 4.4.0, using the packages “car” and “ggpubr”.

#### Protein structure prediction, modeling, and docking

Predictions of protein structures were created with AlphaFold3 (27), installed locally on High Performance Computing (HPC) facility ALICE. Structures were predicted using 100 randomly generated seeds, from which the highest confidence scoring models were selected by the chain pair predicted aligned error (PAE) values. Visualization was performed using PyMol to extract interacting residues.

#### Sequence motif analysis

To identify conserved motifs, the Multiple Expectation Maximization for Motif Elicitation (MEME) software suite (28) was used with the MEME algorithm (version 5.3.2) (29) to discover novel, ungapped motifs without indels. Default settings were used with the anr (any number of repeats) modus, and the maximum number of motifs was set to 8. From the phylum Actinomycetota, DivIVA proteins from the following species were used: Order Mycobacteriales: *Corynebacterium camporealensis, C. glutamicum, M. smegmatis, Mycobacterium tuberculosis, M. xenopi, Rhodococcus erythropolis*; Order Micrococcales: *Micrococcus luteus*; Order Streptomycetales: *K. viridifaciens, Streptomyces coelicolor, S. hygroscopicus*. From the phylum Bacillota, DivIVA proteins from the following species were used: Order Bacillales: *Bacillus atrophaeus, B. pseudomycoides, B. subtilis*; Order Eubacteriales: *Clostridium difficile*; Order Lactobacillales: *Enterococcus faecalis, E. villorum, Streptococcus pneumoniae, S. pyogenes, S. suis*; Order Paenibacillales: *Paenibacillus mucilaginosus*.

## Results

### Construction and functional analysis of DivIVA variants

DivIVA is essential for polar growth of Actinomycetota, but it can be deleted in *Kitasatospora viridifaciens* L-forms, as these replicate via biophysical processes instead of hyphal extension (18). Strain Δ*divIVA* cannot revert to mycelial growth, except if it is complemented by a functional copy of *divIVA* (18). In streptomycetes, DivIVA consists of five domains: an N-terminal domain, followed by a smaller coiled-coil, an intercoil region, a larger coiled-coil, and a C-terminal domain (5). Alphafold3 (AF3) models predict that the intercoil domain and C-terminal domain of *K. viridifaciens* DivIVA are unstructured, as shown by the PAE plot (Fig. 1A). To determine which of these five domains are essential for reversion, we engineered *divIVA* alleles encoding variants of these five domains (Fig. 1B, C) and expressed them in the *divIVA* null mutant. All generated mutant strains expressing one of the DivIVA variants retained their capacity to grow as L-forms (Fig. 1B). However, transferring the strains to reversion plates (21) revealed major differences in terms of their morphology and their efficiency in reverting to a walled state. As expected, expression of wild-type DivIVA enabled the Δ*divIVA* mutant to revert (Fig. 1B). Reversion could not be achieved when the N-terminal domain was absent (Fig. 1B), consistent with previous studies (30) (4). Notably, reversion occurred not only when either the intercoil region or the C-terminus of DivIVA were removed (Fig. 1B), but also when both of these unstructured domains were deleted (Fig. 1B). Deletion of one or both of the unstructured domains resulted in significantly smaller and asymmetrical colonies as compared to strains expressing the wild-type protein, with notable differences in roundness calculated by solidity (pairwise t-test with Holm multiple testing correction, p < 0.0001) (Figs. 1D, 1E). Furthermore, we detected no significant difference in the reversion efficiency among the complemented strains. Still, all these strains that expressed *divIVA in trans* reverted significantly less efficiently than *alpha* (one-way ANOVA, p < 0.0001) (Fig. 1F). When additionally the second coiled-coil domain of DivIVA was deleted from the version already lacking the intercoil region or the C-terminal part, the ability to complement Δ*divIVA* and support reversion was lost (Fig. 1B), indicating that DivIVA_ΔIR/C_, hereinafter referred to as DivIVA_min_, represents a minimized DivIVA protein capable of supporting walled growth in *K. viridifaciens*.

**Figure 1.**
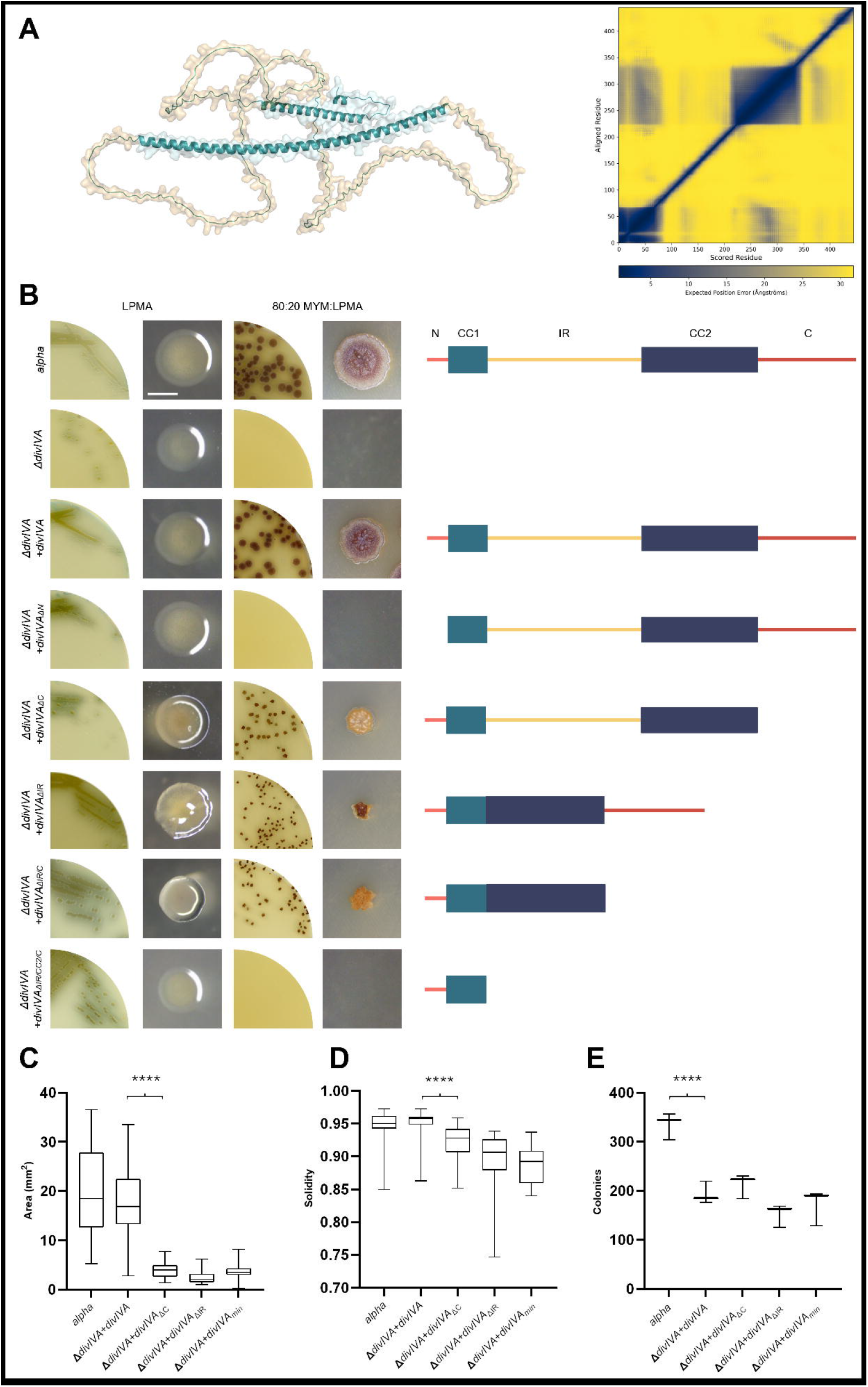
Phenotypic analysis of *K. viridifaciens* Δ*divIVA* expressing *divIVA* variants. A) Cartoon and surface representation of DivIVA of *K. viridifaciens* made using AF3. Structured domains are blue, and unstructured domains are light brown. The PAE plot is shown on the right. Low confidence values are yellow, high values are blue. pLDDT values - per-residue measures of local confidence - can be found in Figure S1. B) L-form strains expressing different *divIVA* alleles were tested for growth as L-forms on LPMA medium (left two images) and for the ability to revert to walled growth on non-osmoprotective plates (right two images). Scalebar = 2 mm. C) Schematic overviews showing the architecture of the DivIVA variants from panel A, relative to the wild-type protein (top), which contains an N-terminal region (N) (light red), a smaller coiled-coil (CC1) (light blue), an intercoil region (IR) (yellow), a larger coiled-coil (CC2) (dark blue), and a C-terminal domain (C) (dark red). D) Quantification of the average colony area for strains expressing DivIVA variants (N=30). E) Quantification of colony solidity for strains expressing DivIVA variants (N=30). F) The number of reverted colonies per strain per reversion plate (N=3).

### Deletion of the unstructured domains affects mycelial morphology

To further study the effect of deleting the unstructured domains of DivIVA on growth and morphogenesis, reverted colonies were inoculated in TSBS medium and subsequently transferred to TSBS medium in non-shaking flasks. As expected, the morphology of the strain expressing the ectopic copy of native *divIVA* was identical to that of *alpha* (Fig. 2A). Deleting either the intercoil or the C-terminal region impacted branching dynamics, as mycelia of Δ*divIVA+divIVA_ΔIR_* and Δ*divIVA+divIVA_ΔC_* were irregular and more highly branched with branches closer to the tip compared to Δ*divIVA+divIVA* (Fig. 2A). Δ*divIVA+divIVA_min_* showed a more extreme phenotype characterized by short and wide hyphae of highly irregular shape and cell dimension, and more compact mycelial clumps than in the single domain deletions (Fig. 2A). Live-cell time-lapse imaging showed that, in contrast to *ΔdivIVA+divIVA* (Video S1), hyphae of Δ*divIVA+divIVA_min_* showed frequent localized bursting of hyphae at the tips (Fig. 2B, Video S2-3). Scanning electron microscopy revealed that hyphae of *ΔdivIVA+divIVA_min_* exhibited an abnormal, widened morphology, characterized by kinked shapes and short outgrowths (Fig. 2C). Measurements of hyphal thickness in scanning electron micrographs revealed that the hyphae were, on average, 1.7x wider in *ΔdivIVA+divIVA_min_* compared to *ΔdivIVA+divIVA* (Fig. 2F). Transmission electron microscopy revealed an average 1.3 x increase in the number of septa in *ΔdivIVA+divIVA_min_* compared to *ΔdivIVA+divIVA* (N=20) (Fig. 2D and 2G). Using cryo-electron tomography, we also found an average 1.9 x increase in subapical cell wall thickness between *ΔdivIVA+divIVA* and *ΔdivIVA+divIVA_min_* (Fig. 2E and 2H). In conclusion, these data show that deletions of the unstructured domains, namely the intercoil region and C-terminus, have pleiotropic effects on the morphology, affecting branching frequency, and that deleting both these domains to make the *divIVA_min_* allele leads to a very strong phenotype, with increasing cross-wall formation, highly irregular hyphal diameter, increased cell-wall thickness, and frequent lytic events.

**Figure 2.**
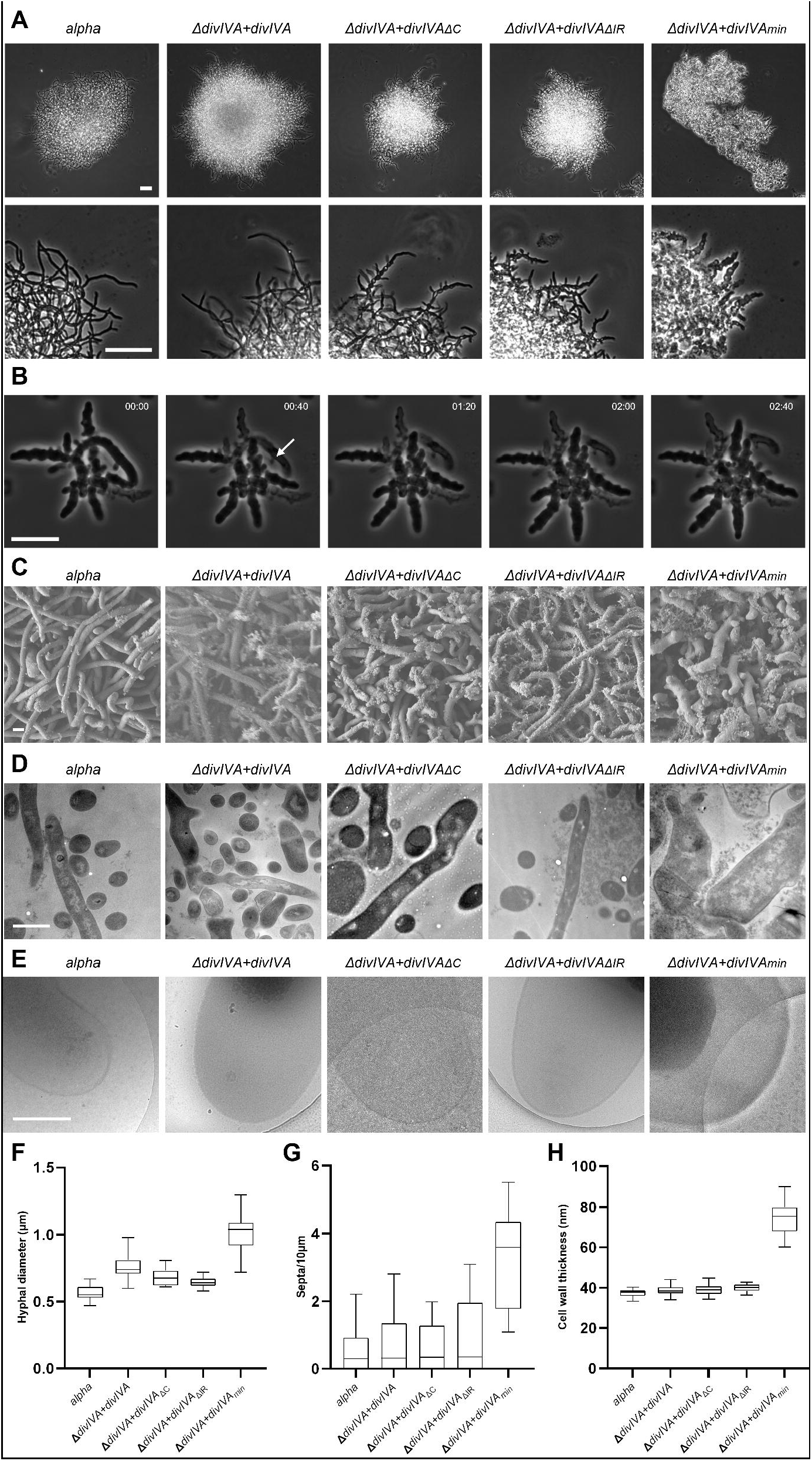
Morphological characterization of strains expressing distinct DivIVA variants. A) Phase contrast microscopy pictures of mycelia of Δ*divIVA* transformants expressing different *divIVA* variants grown overnight in TSBS without shaking. Scalebar = 20 µm. B) Time-lapse sequence showing bursting of a hypha (white arrow) of *ΔdivIVA+divIVA_min_* observed in a microfluidics chamber. Mycelia were grown in TSBS and loaded into a CellASIC ONIX2 system. Time is in hr:min. Scalebar = 10 µm. C) Scanning electron micrographs of strains *Alpha* and *ΔdivIVA+divIVA*, ΔdivIVA+divIVA_ΔC_, ΔdivIVA+divIVA_ΔIR,_ and *ΔdivIVA+divIVA_min_* grown as mycelium on MYM:LPMA (4:1). Scalebar = 1 µm. D) Transmission electron micrographs of thin-sections of mycelia of the five revertant strains grown on MYM:LPMA (4:1). 70 nm slices were made and placed on copper TEM support grids. Scalebar = 1 µm. E) Cryo-electron micrographs of hyphal tips of five revertant strains. Isolated sacculi were vitrified and observed inside a 120 kV Talos L120C TEM. Scalebar = 500 nm. F) Hyphal diameter of the five revertant strains calculated from SEM data (N=15). G) The number of septa measured for mycelia with a length >4 µm calculated from TEM data (N=12). H) The average subapical cell wall thickness measured for each strain calculated from cryo-electron micrographs (N=15). Cell wall thickness was calculated by averaging pixel values (arbitrary units) alongside the straightened cell wall and normalized against the background pixel values.

### Chimeric DivIVA functionality in K. viridifaciens is limited to Actinomycetota

AF3 modeling predicts that deletion of the intercoil region and the C-terminal domain of DivIVA results in a fusion of the two coiled-coil domains (Fig. 3A). Furthermore, predicted structures of DivIVA_min_ revealed that its self-association results from a C-terminal motif that enables antiparallel dimer-dimer binding, resulting in a tetramer (Fig. 3A). Despite superficial structural similarity with *B. subtilis* DivIVA (Fig. 3A), residue-level structural comparisons reveal that *B. subtilis* DivIVA and DivIVA*min* differ significantly in the amino acids responsible for the homodimerization and subsequent dimer-dimer tetramerization. AF3 models suggest that homodimers are primarily formed through self-binding in the N-terminus, with a weak interaction between the CC1 domains of monomers. The CC2 domain notably only becomes important in dimer-dimer interactions between two antiparallel homodimers (Fig. 3B). The pLDDT values - per-residue measures of local confidence - of the *B. subtilis* DivIVA tetramers are lower than those of DivIVA_min_ possibly explained by a lower likelihood to tetramerize (Fig. S2). Furthermore, there are sequence differences in motifs found in the first and second coiled-coil domain between Actinomycetota and Bacillota (Table S4). These differences likely reflect the distinct growth modes: *B. subtilis* elongates through lateral insertion of cell wall material (11), whereas *K. viridifaciens* grows by polar extension (31). Furthermore, Alphafold3 (AF3) models predict that DivIVA from *M. smegmatis* has an unstructured region starting at the same position as in *K. viridifaciens* DivIVA (Fig. 3A, Fig. S3).

**Figure 3.**
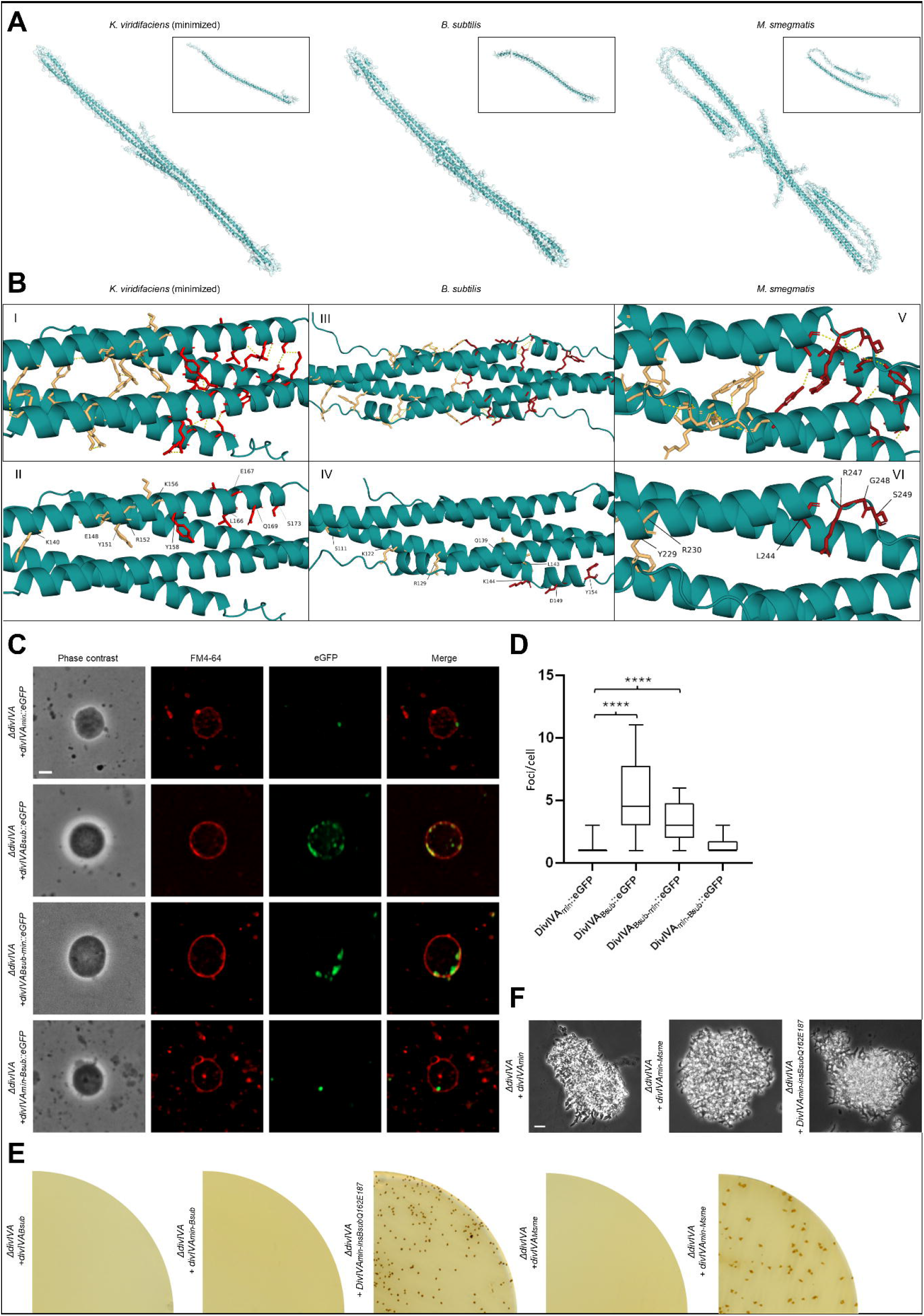
Chimeric DivIVA of *Mycolicibacterium smegmatis* can facilitate reversion. A) Cartoon and surface representation of monomers and predicted tetramers of minimized DivIVA of *K. viridifaciens*, and DivIVA from *B. subtilis* and *M. smegmatis*, made using AF3. PAE plots can be found in Figure 3. B) Residue-level models of the putative tetramerization domain in minimized DivIVA *of K. viridifaciens*, and DivIVA from *B. subtilis* and *M. smegmatis*. The two sides of the antiparallel binding are colored light orange and red. C) eGFP was fused to the C-terminus of various DivIVA proteins to check for self-association behavior. Image stacks along the Z-axis (spacing 270 nm) were collected and deconvolved, and orthogonal projections were made from the four focal planes around the center of the cells. L-form cells were placed on 1% agarose in P-buffer with 0.5 μg mL^-1^ FM4-64 for membrane labeling. Scalebar = 2 µm. D) The number of DivIVA::eGFP foci was counted per cell throughout the orthogonal projections (N=20). E) L-form strains expressing different chimeric *divIVA* alleles were tested for the ability to revert to walled growth on non-osmoprotective plates. F) Phase contrast microscope pictures of mycelia of Δ*divIVA+DivIVA_min-insBsubQ162E187_* and Δ*divIVA+divIVA_min-Msme_* grown overnight in TSBS without shaking. Scalebar = 10 µm.

Complementation experiments revealed that *B. subtilis* DivIVA could not restore walled growth in the *ΔdivIVA* mutant of *K. viridifaciens* (Fig. 3E). Localization studies using C-terminal eGFP fusions showed that this failure was likely due to improper localization: *B. subtilis* DivIVA formed multiple foci instead of a single, well-defined focus (Fig. 3C). Replacing its N-terminal domain with that of DivIVA_min_ significantly improved localization, resulting in a single focus similar to DivIVA_min_ (Fig. 3C). Still, this improvement was not sufficient to restore functionality (Fig. 3E). Similarly, swapping the C-terminal region of *B. subtilis* DivIVA with that of DivIVA_min_ also failed to support reversion. Interestingly, functionality was maintained when only the last 26 amino acids of DivIVA_min_ were replaced with those from *B. subtilis* (Fig. 3E-F). This likely reflects the limited role of these residues as part of an unstructured tail in the antiparallel binding structure downstream in the second coiled-coil region (Fig. 3B).

By contrast, while native DivIVA from the polar-growing *M. smegmatis* could not restore reversion, a chimeric *M. smegmatis* DivIVA protein in which the N-terminal domain was replaced with that of *K. viridifaciens* could (Fig. 3E), resulting in a morphology similar to that of DivIVA_min_ (Fig. 3F). Together, these results suggest that, despite structural similarities, DivIVA proteins have diverged functionally in response to different growth modes. This highlights the importance of specific amino acid sequence elements for proper localization, differentiation, and cellular function.

## Discussion

DivIVA is a coiled-coil protein widely conserved in both Actinomycetota and Bacillota. In Actinomycetota, such as *K. viridifaciens* and *M. smegmatis*, DivIVA plays an essential role in polar growth (5, 6, 8), whereas in Bacillota, such as *B. subtilis*, DivIVA plays a non-essential role in cell division (11). So far, it has been impossible to functionally replace *divIVA* in Streptomycetaceae with those of other taxa. To overcome this limitation, we here exploited an L-form *divIVA* deletion strain derived from *K. viridifaciens*, as this strain replicates via biophysical processes instead of hyphal extension (18). These cells can only reinitiate walled growth in the presence of a functional form of DivIVA. Using this platform, we have investigated the functionality of truncated DivIVA proteins. After deleting the intercoil region and the C-terminal domain, both of which we have shown to be unstructured domains, we have built a minimized DivIVA protein consisting of only the N-terminus and two fused coiled-coils.

DivIVA forms a multimeric complex in streptomycetes and long filaments *in vitro* (4). Our AlphaFold3 models predict that DivIVA_min_ assembles into a tetramer, similar to DivIVA of *B. subtilis* (13). When we deleted the second coiled-coil domain in DivIVA_min_, this hampered reversion to a walled state in *ΔdivIVA*. In *Streptomyces coelicolor*, deleting the second coiled-coil of DivIVA disrupted oligomerization (4). In *Corynebacterium glutamicum*, the N-terminal domain and two coiled-coil regions have already been shown to be essential for DivIVA functioning (30). In *B. subtilis*, the C-terminal coiled-coil region, which correlates to the second coiled-coil in *K. viridifaciens*, is involved in oligomerization (13). In *Deinococcus radiodurans*, purified DivIVA showed bundles under TEM, whereas the N-terminal variant, including the coiled-coil, did not (32). Furthermore, the C-terminal region seemed to be crucial for the structural and functional integrity of DivIVA (32). Altogether, this shows that the multimerization behavior of DivIVA is conserved amongst bacterial taxa.

Although our minimal DivIVA protein is structurally similar to that of *B. subtilis*, complementation of *B. subtilis* DivIVA in Δ*divIVA* could not facilitate reversion to a walled state. Substituting the N-terminal domain with that of *K. viridifaciens* resulted in a comparable number of foci to DivIVAmin, yet still failed to support reversion. This may reflect differences in membrane affinity, as simulations have predicted distinct membrane interactions for DivIVA from *S. coelicolor* and *B. subtilis* (10). Notably, Actinomycetota possess different amino acid motifs in their coiled-coil domains compared to Bacillota, suggesting functional divergence. Previous studies have shown that DivIVA from *Mycobacterium tuberculosis* and *Streptomyces coelicolor* can compensate for DivIVA loss in *C. glutamicum* and restore polar growth, whereas *B. subtilis* DivIVA cannot (7). Interestingly, either coiled-coil domain could be swapped with the corresponding domain from *B. subtilis* without loss of function, provided the N-terminal domain of *C. glutamicum* DivIVA remained intact (30). In our experiments, replacing only the last 26 amino acids of DivIVAmin with those from *B. subtilis* enabled reversion. In contrast, when we substituted the N-terminal domain with that of *K. viridifaciens*, DivIVA from *M. smegmatis* facilitated reversion. Together, these results indicate that polar growth in Streptomycetaceae depends not only on overall structural features but also critically on specific amino acid sequences.

Deleting either the intercoil or the C-terminal region impacted branching dynamics, resulting in irregularly shaped strains with increased branching and shorter distances between the tips and new branches. Deletion of both domains caused more severe morphological defects, including irregular hyphal diameter, reduced septation, thinner cell walls, and frequent tip bursting. These defects may result from the loss of interactions with key partner proteins. DivIVA is known to interact with several cytoskeletal components, including Scy and FilP, forming a dynamic complex at the hyphal tip that coordinates the insertion of new cell wall material (33-35). It also interacts with CslA, a cellulose synthase-like protein involved in β-glucan biosynthesis at growing tips (36-38). In our study, deletion of the C-terminal region likely disrupted DivIVA phosphorylation, as five *in vivo* phosphorylation sites have been identified in this region in *Streptomyces coelicolor* (39). One of the kinases responsible is AfsK, a serine/threonine protein kinase (40), while the phosphatase SppA reverses this modification (23). Future studies should focus on clarifying the roles of the unstructured regions and identifying which specific interaction partners are lost upon their deletion. In conclusion, our findings indicate that while a minimized DivIVA protein can initiate polar growth in Streptomycetaceae, it is insufficient to maintain the typical mycelial architecture. This highlights a crucial, yet undefined role for the unstructured regions in the morphogenesis of Streptomycetaceae.

## Acknowledgments

We thank Gerda Lamers for assisting with the scanning electron microscope and Le Zhang for constructive discussions. We are also grateful to Lennart Schada von Borzyskowski for providing us with the *E. coli* TOP10 strain. We also thank Duschka Kleijn for her feedback on the manuscript.

Maarten Lubbers: Conceptualization, Data curation, Formal analysis, Investigation, Methodology, Visualization, Writing. Belmin Bajramović: Formal analysis, Investigation, Methodology, Visualization, Writing. Véronique Ongenae: Investigation, Methodology, Visualization. Joost Willemse: Methodology, Visualization. Dieuwertje de Bruin: Formal analysis, Investigation, Methodology. Niels Mulder: Investigation, Methodology. Bastienne Vriesendorp: Methodology. Francisco Barona-Gómez: Methodology, Writing – review & editing. Ariane Briegel: Methodology, Writing – review & editing. Klas Flärdh: Methodology, Resources, Writing – review & editing. Gilles P. van Wezel: Methodology, Resources, Supervision, Writing – review & editing. Dennis Claessen: Conceptualization, Funding acquisition, Project administration, Resources, Supervision, Writing – review & editing.

## Supplementary data

**Figure S1.**
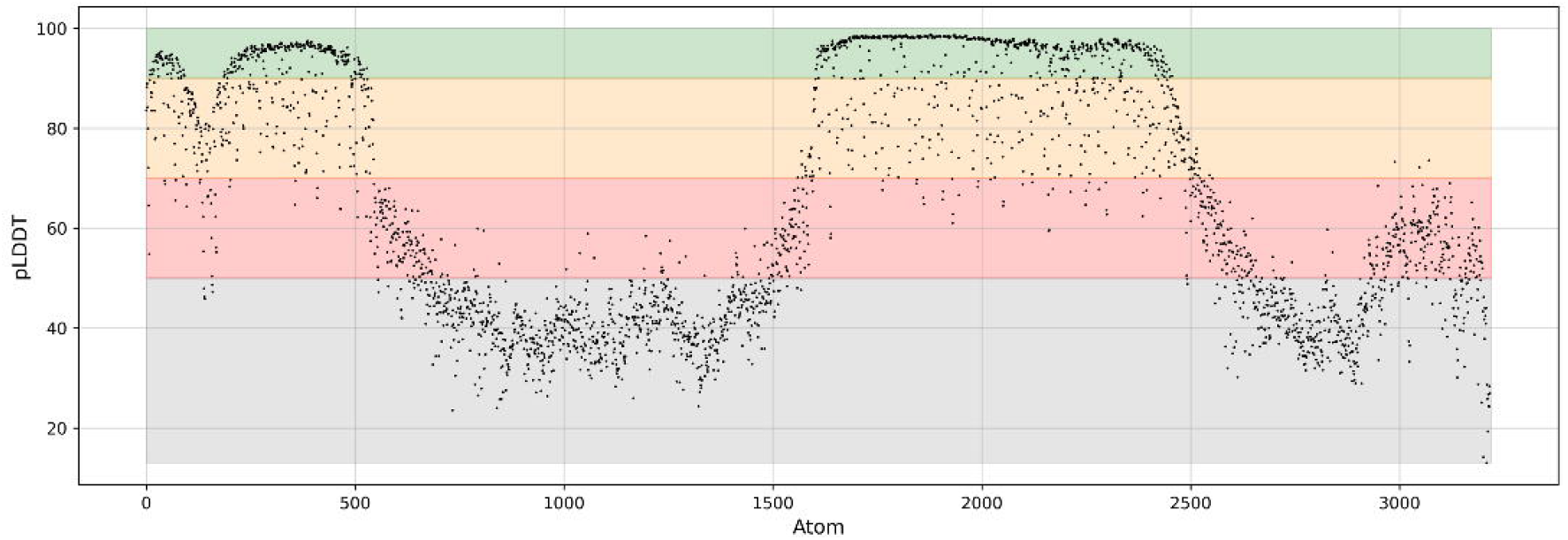
Atomic-level pLDDT values of DivIVA of *K. viridifaciens*. Confidence levels are given by four different colors: Green: very high (90-100); yellow: high (70-90); red: low (50-70); grey: very low (<50).

**Figure S2.**
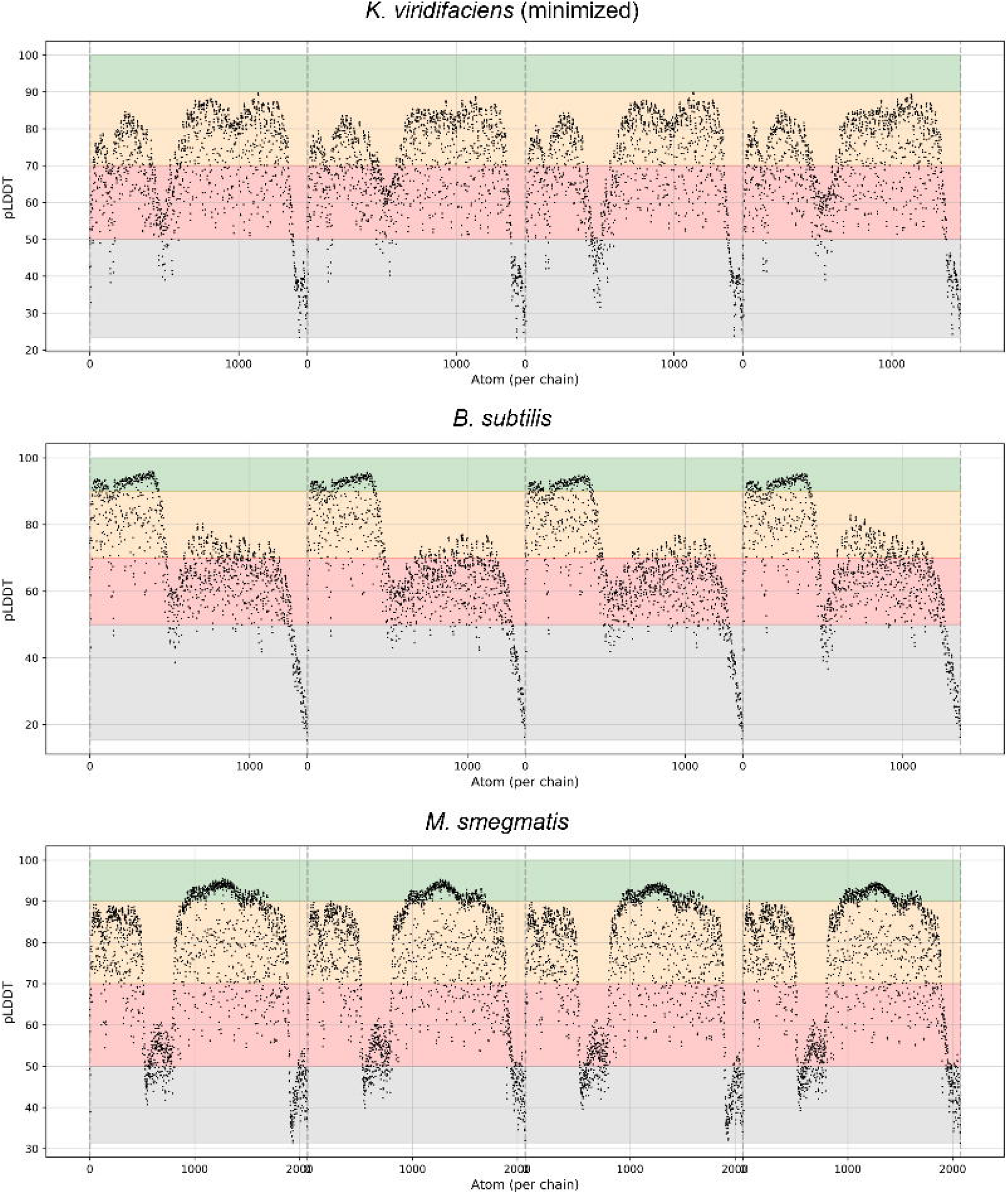
Atomic-level pLDDT values of the predicted tetramers of minimized DivIVA of *K. viridifaciens* (minimized), and the DivIVA proteins of *B. subtilis* and *M. smegmatis*. Confidence levels are given by four different colors: Green: very high (90-100); yellow: high (70-90); red: low (50-70); grey: very low (<50).

**Figure S3.**
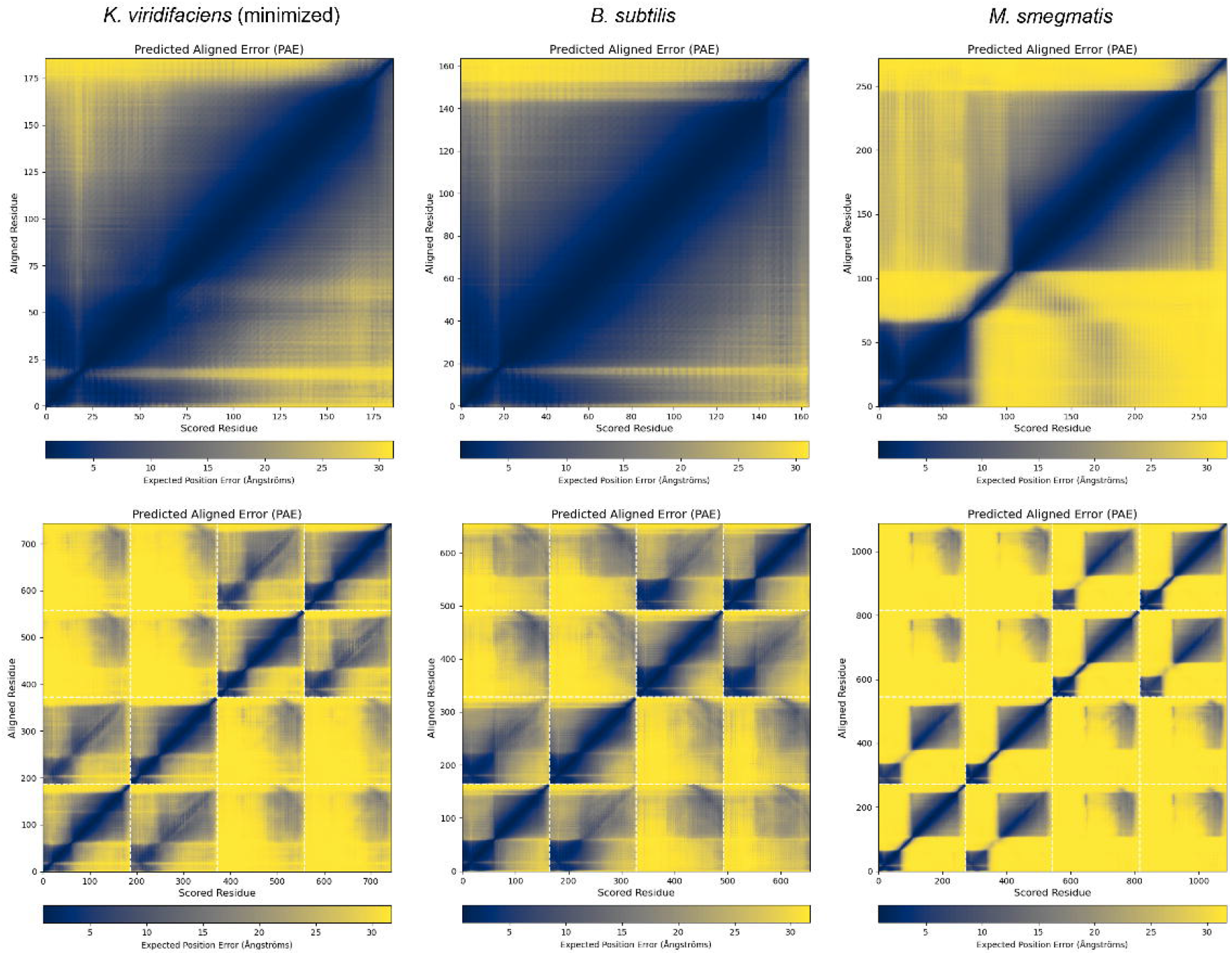
PAE plots of the monomers (top row) and tetramers (bottom row) of minimized DivIVA of *K. viridifaciens*, and the DivIVA proteins of *B. subtilis* and *M. smegmatis*. Low confidence values are given in yellow, high values in blue.

*Video S1:* Growth of Δ*divIVA+divIVA* observed in a growth chamber. MYM medium was enclosed between a membrane and a cover slip and incubated at 30 °C. Scalebar = 10 µm. Time is in min.

*Video S2:* Growth of Δ*divIVA+divIVA*_min_ observed in a growth chamber. MYM medium was enclosed between a membrane and a cover slip and incubated at 30 °C. Scalebar = 10 µm. Time is in min.

*Video S3:* Bursting of hyphae of Δ*divIVA+divIVA*_min_ observed in a CellASIC B04A-03 microfluidics chamber. Scalebar = 10 µm. Time is in min.

**Table S1:**
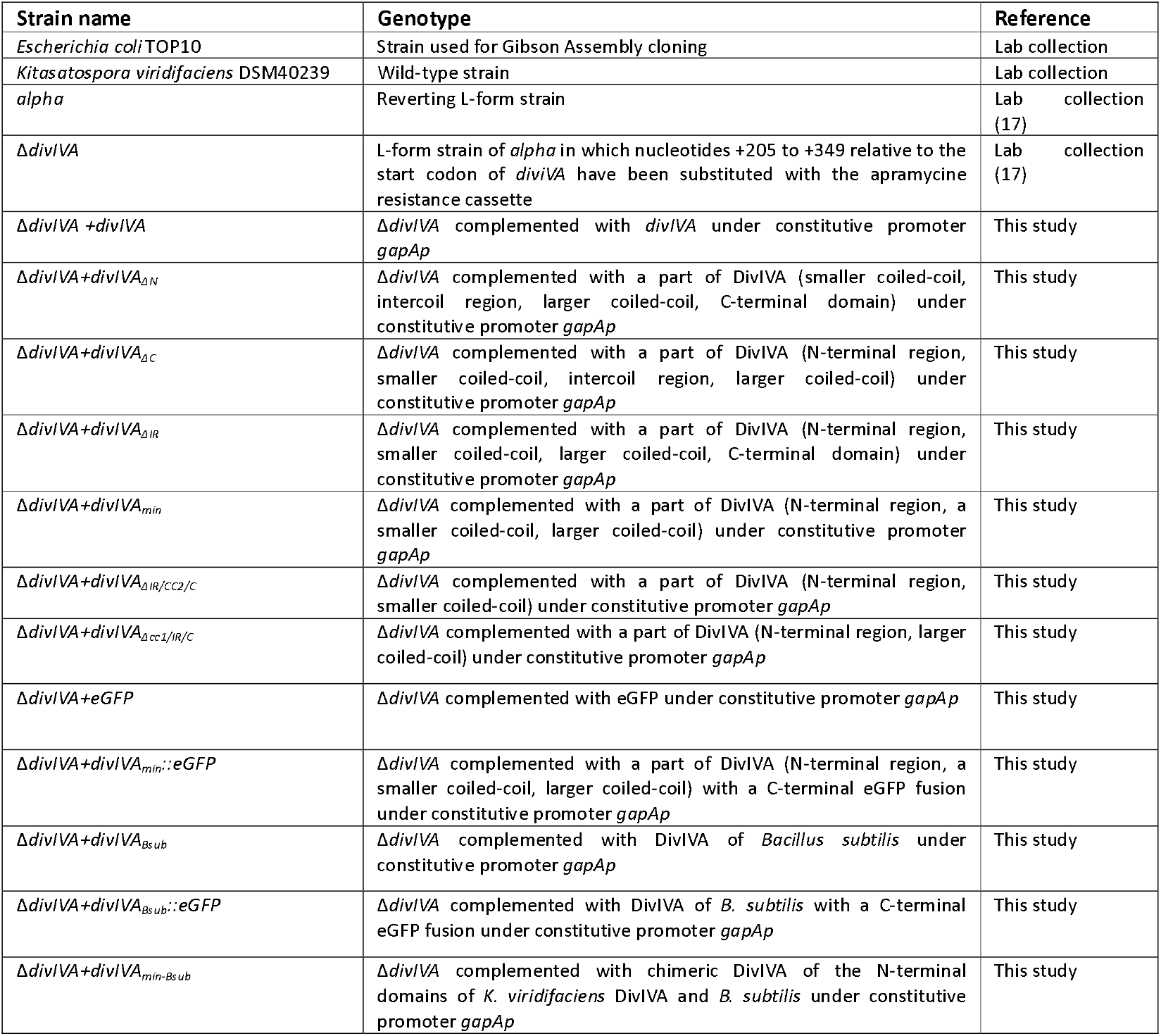
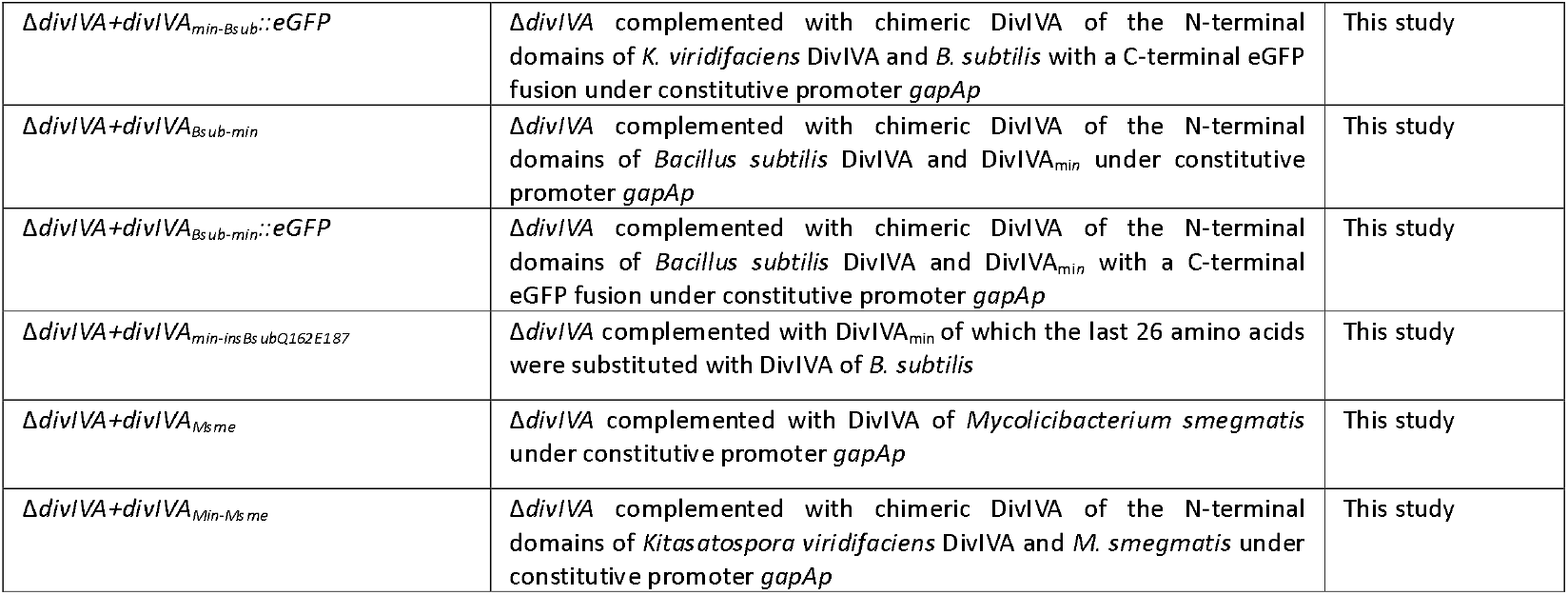
Strains used in this study.

**Table S2:**
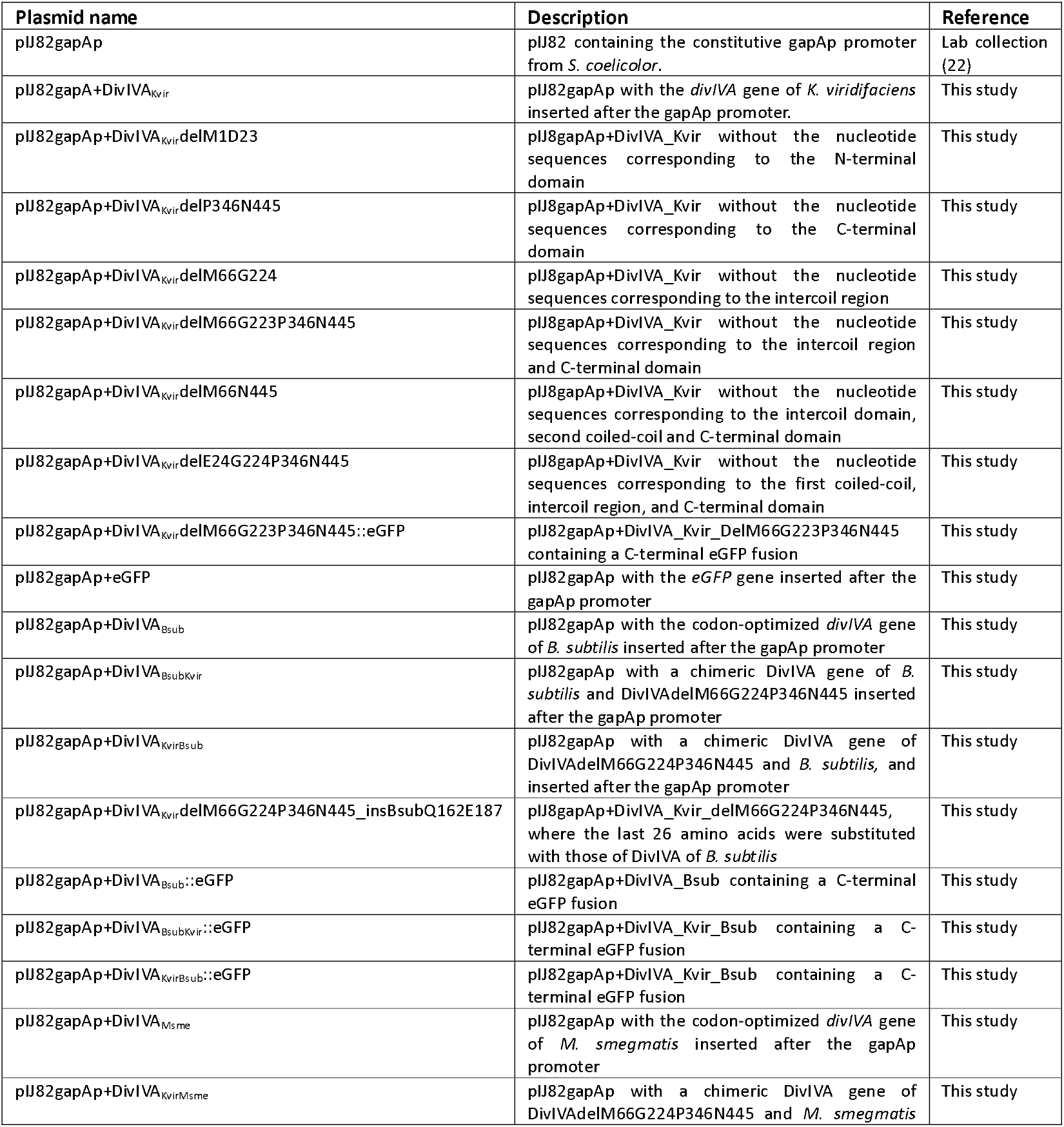
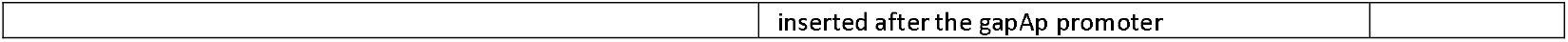
Plasmids used in this study.

**Table S3:**
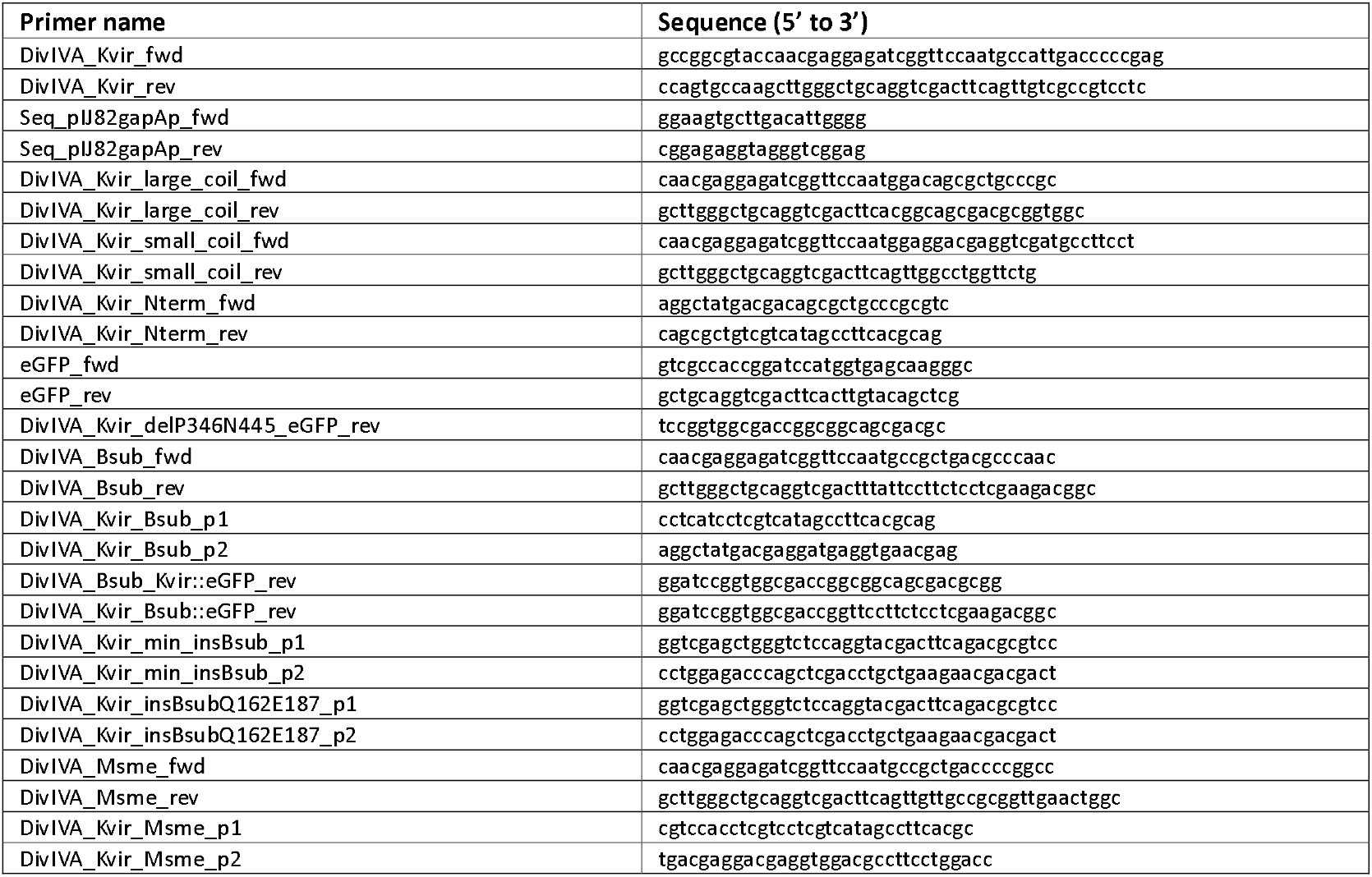
Primers used in this study.

**Table S4:** Motifs present in the first and second coils in Actinomycetota and Bacillota.

| Motif | Position | Phylum | Sequence | Sites | E-value | Width |
| --- | --- | --- | --- | --- | --- | --- |
| 1 | First coiled-coil | Actinomycetota |  | 10 | 1.4e-089 | 41 |
| 1 | First coiled-coil | Bacillota |  | 10 | 1.1e-080 | 41 |
| 2 | Second coiled-coil | Actinomycetota |  | 10 | 1.4e-167 | 50 |
| 2 | Second coiled-coil | Bacillota |  | 9 | 7.8e-102 | 50 |
| 3 | Second coiled-coil | Actinomycetota |  | 15 | 1.1e-094 | 41 |
| 3 | Second coiled-coil | Bacillota |  | 10 | 3.7e-033 | 21 |

